# Manganese-dependent microRNA trimming by 3’→5’ exonucleases generates 14-nucleotide or shorter tiny RNAs

**DOI:** 10.1101/2022.10.06.511180

**Authors:** GeunYoung Sim, Audrey C. Kehling, Mi Seul Park, Jackson Secor, Cameron Divoky, Huaqun Zhang, Nipun Malhotra, Divyaa Bhagdikar, Ekram Abd El-Wahaband, Kotaro Nakanishi

## Abstract

MicroRNAs (miRNAs) are about 22-nucleotide (nt) non-coding RNAs forming the effector complexes with Argonaute (AGO) proteins to repress gene expression. Although tiny RNAs (tyRNAs) shorter than 19 nt have been found to bind to plant and vertebrate AGOs, their biogenesis remains a long-standing question. Here, our *in vivo* and *in vitro* studies show several 3’→5’ exonucleases, such as interferon-stimulated gene 20 kDa (ISG20), three prime repair exonuclease 1 (TREX1), and ERI1 (enhanced RNAi, also known as 3’hExo), capable of trimming AGO-associated full-length miRNAs to 14 nt or shorter tyRNAs. Their guide trimming occurs in a manganese-dependent manner but independently of the guide sequence and the loaded four human AGO paralogs. We also show that ISG20-mediated guide trimming makes Argonaute3 (AGO3) a slicer. Given the high Mn^2+^ concentrations in stressed cells, virus-infected cells, and neurodegeneration, our study sheds light on the roles of the Mn^2+^-dependent exonucleases in remodeling gene silencing.

## Introduction

In the canonical miRNA biogenesis, Dicer processes the hairpin-structured precursor miRNAs into about 22-nt miRNA duplexes (1). AGOs load the duplexes and eject the passenger strand, forming the effector complexes of the RNA interference (RNAi) (2, 3). Previous RNA sequencing studies identified AGO-associated 10–18-nt tyRNAs in plants and vertebrates that mapped onto tRNAs and miRNAs (4). However, how those tyRNAs are synthesized remains unknown. The present study focused on the biogenesis of miRNA-derived tyRNAs.

## Results

We raised several hypothetical pathways towards AGO-associated tyRNAs (**Fig. 1A**). If short single-stranded (ss) RNAs are directly loaded into AGOs, they must exist stably in the cell. To test the idea, we transfected 14- or 23-nt ss miR-20a or their small interfering RNA (siRNA)-like duplex into HEK293T cells. Only the guide RNA was radiolabeled at its 5’ end. Both 14- and 23-nt ssRNAs and the 14-nt siRNA-like duplex were degraded (**Fig. 1B**), suggesting that neither has a chance of being loaded into AGOs efficiently. In contrast, the 23-nt siRNA-like duplex remained 19-23 nt, and no tyRNA was detected, indicating that a detectable level of tyRNAs is not generated in normal conditions. These results prompted us to think that after the canonical RISC assembly, the AGO-associated miRNAs are trimmed to tyRNAs by yet unidentified 3’→5’ exonucleases (**Fig. 1A bottom pathway**). Previous studies reported the expression of ISG20, a 3’→5’ exonuclease, induced by interferon upon viral infection and stress and by estrogen hormone (5, 6). When ISG20 was exogenously expressed, a small amount of 13-14 nt was detected (**Fig. 1C**). To test whether the generated tyRNA is associated with AGO, FLAG-AGO2 was co-expressed with ISG20 in HEK293T cells (**Fig. 1D bottom**) that were also transfected with an siRNA duplex whose 23-nt miR-20a guide is radiolabeled at its 5’ end. After 48 hours, FLAG-AGO2 was immunopurified, and the associated RNAs were resolved on a denaturing gel. Co-expression of ISG20 generated AGO-associated tyRNAs (**Fig. 1D top**). tyRNA generation was also observed when TREX1 or ERI1 was co-expressed instead of ISG20. ERI1 is known to negatively regulate global miRNAs abundance in mouse lymphocytes (7), but the mechanism remains unclear. Poly(A)-specific ribonuclease (PARN) shortened the 23 nt guide down to 19 nt, whereas exonuclease 5 (EXO5) did not trim the guide at all. The experiment was repeated with their catalytically dead mutants to confirm that the observed trimming was due to their exonuclease activity. As a result, none of the mutants generated tyRNA **(Fig. 1D)**, proving that the catalytic center of the exonucleases is essential for tyRNA generation. ISG20, TREX1, and ERI1 also generated tyRNAs from AGO1-, AGO3-, and AGO4-associated miR-20a (**Fig. 1E**).

**Figure 1.**
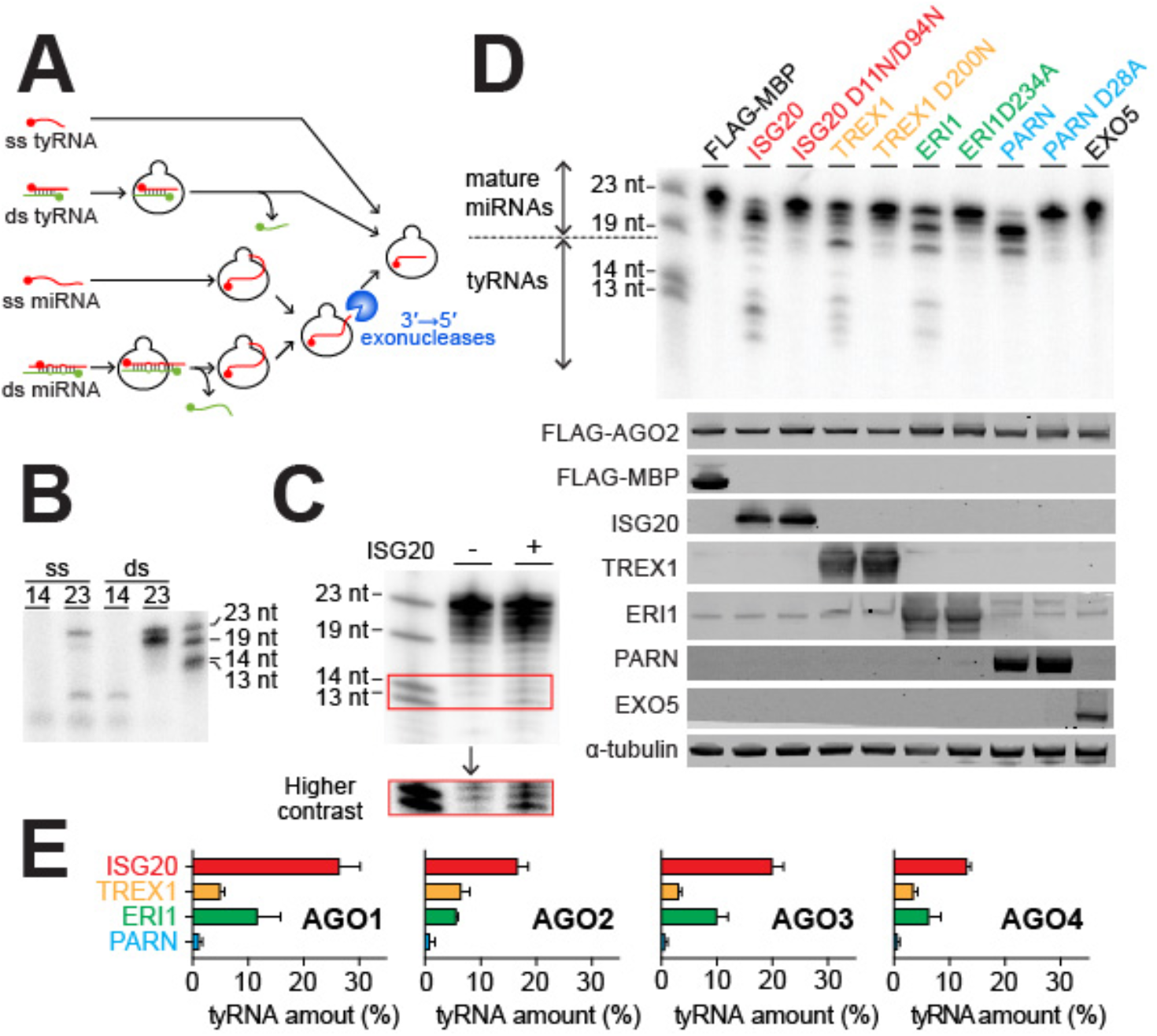
Specific 3’→5’ exonucleases convert miRNAs to tyRNAs. (*A*) Possible pathways towards AGO-associated tyRNAs. (*B*) *In vivo* stability of 14- and 23-nt ss miR-20a and their siRNA-like duplexes (*C*) Accumulation of tyRNAs upon expression of ISG20. (*D*) *In vivo* trimming of FLAG-AGO2-associated miR-20a by ISG20, TREX1, ERI1, PARN, and EXO5 and their catalytic mutants. A representative gel image (top) and western blots with antibodies for each protein (bottom). (*E*) tyRNA synthesis on four human AGOs by ISG20, TREX1, ERI1, and PARN.

To characterize their guide trimming, we reconstituted an *in vitro* trimming system (see SI). FLAG-tagged human AGO1, AGO2, AGO3, and AGO4 were programmed with a 5’-end radiolabeled 23-nt miR-20a, immobilized on anti-FLAG beads to wash out free guide RNAs, and incubated with one of the abovementioned exonucleases in the presence of manganese. All exonucleases, except for EXO5, trimmed miR-20a that had been loaded onto slicer-deficient AGO1 and AGO4 as well as slicing-competent AGO2 and AGO3 (8) (**Fig. 2A left**). In contrast, the catalytic mutants of the exonucleases showed no trimming activity (**Fig. 2B**), further supporting our conclusion that the observed guide trimmings are solely due to the catalytic activity of the exonucleases, not that of the AGOs. Each exonuclease similarly trimmed the loaded miR-20a across the four AGOs, but the trimming patterns differed between the exonucleases (**Fig. 2A right**). PARN generated an abundance of 21 nt. ERI1 ceased trimming when the guide length became 19 nt while developing a small population of 14-15-nt tyRNAs. In contrast, ISG20 and TREX1 shortened miR-20a to 13 and 14 nt, respectively. Replacing manganese with magnesium drastically reduced the trimming activities of ISG20, TREX1, and ERI1, but not PARN (**Fig. 2C**). These results suggest that these exonucleases can synthesize tyRNAs from miRNAs in a Mn^2+^-dependent manner with different processing rates.

**Figure 2.**
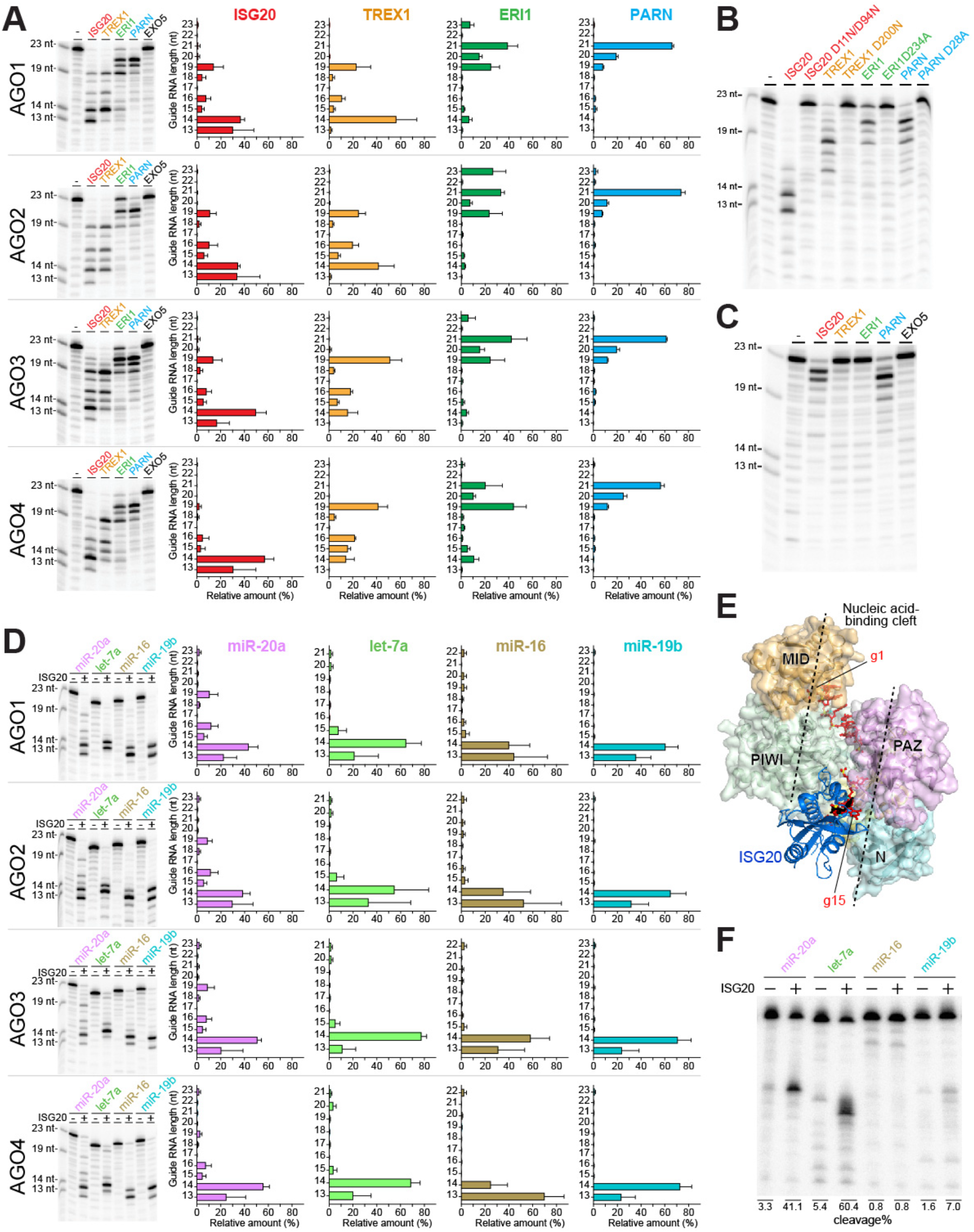
ISG20, TREX1, and ERI1 generate tyRNAs autonomously. (*A*) *In vitro* guide trimming by 3’→5’ exonucleases in the presence of 2 mM MnCl_2_. (*Left*) A representative gel image for each AGO. (*Right*) The relative amount of each guide length after incubation with either 3’→5’ exonuclease. (*B*) *In vitro* trimming of AGO2-associated miR-20a by the catalytically dead exonuclease mutants at 2 mM MnCl_2_. (*C*) *In vitro* trimming of AGO2-associated miR-20a by 3’→5’ exonucleases in 2 mM MgCl_2_. (*D*) *In vitro* trimming of different miRNAs by ISG20. (*Left*) A representative gel image for each AGO. (*Right*) The relative amount of each guide length after incubation with ISG20. (*E*) Docking model of ISG20 (blue ribbon model) on a guide (red stick model)-bound AGO3 (surface model). (*F*) *In vitro* trimming of AGO3-associated different miRNAs, followed by target cleavage.

Next, we investigated the susceptivity of different miRNAs to the 3’→5’ exonucleases. To this end, FLAG-tagged AGOs were programmed with a 5’-end radiolabeled 23-nt miR-20a, 21-nt let-7a, 22-nt miR-16, or 23-nt miR-19b, followed by incubation with ISG20. Although miR-20a was trimmed slower than the others, all the tested miRNAs were shortened to 13-14 nt, regardless of which AGO was loaded (**Fig. 2D**). This suggests that ISG20 can generate tyRNAs from a variety of miRNAs that are incorporated into any AGO. Our docking model indicates that the nucleic acid-binding cleft of AGO sequesters the guide nucleotide 1-14 (g1-g14) positions while the remaining 3’ half is accessible to ISG20 (**Fig. 2E**). This could explain why trimming stops when the guide length becomes 13–14 nt. Our previous study revealed that 14-nt variants of miR-20a and let-7a, but not of miR-16 or miR-19b, whose 3’ 7-9 nt were trimmed from their mature miRNAs, conferred a decent slicing activity on AGO3 (8). These results prompted us to test whether trimming of AGO3-associated specific miRNAs makes the RISCs a slicer. To this end, FLAG-AGO3 was programmed with the same full-length miRNAs used in Fig. 2D, incubated with ISG20, and mixed with a cap-labeled target RNA containing a complementary sequence to each guide. Cleavage was observed only when FLAG-AGO3 was programmed with miR-20a and let-7a (**Fig. 2F**), demonstrating that ISG20 catalytically activates AGO3.

## Discussion

ISG20, TREX1, and ERI1 convert AGO-associated miRNAs to tyRNAs, which requires Mn^2+^, an essential transition metal for human health. Dysregulation of the cellular Mn^2+^ concentration has been implicated in neurodegenerative diseases, such as Parkinson’s disease, Alzheimer’s disease, Huntington’s disease, and manganism (9). Notably, the ISG20 level is elevated in neurodegenerative disease models (10) and brain injury (5), while the malfunction of TREX1 causes autoimmune diseases such as Aicardi-Goutieres syndrome (11). Natural killer cells and T cells deficient in ERI1 enhance the RNAi (7). These results suggest a possible correlation between the Mn^2+^-dependent guide-trimming and neurodegenerative diseases.

## Materials and Methods

### Cloning, expression, and purification of recombinant proteins

Recombinant AGOs were purified from insect cells as previously reported (12, 13). Exonucleases, including ISG20, ISG20 D11N/D94N, TREX1, TREX1 D200N, ERI1, ERI1 D234A, PARN, PARN D28A, or EXO5, were cloned into a SUMO-fused pRSFDuet-1 vector (Novagen). Catalytic mutants were created from these constructs with site-directed mutagenesis, using PCR primers designed to anneal to the mutation site. PCR products were subsequently digested by DpnI (NEB) and transformed into DH5α *E. coli* cells (Invitrogen). All plasmids were verified by Sanger sequencing (OSU Shared Resources). Exonuclease plasmids were transformed into BL21 Rosetta 2(DE3) pLysS *E.coli* cells (Novagen) for expression. Cells were harvested by centrifugation at 5,000 xg for 10 minutes at 4°C, then the remaining pellet was resuspended with Lysis Buffer (1x PBS, 500 mM NaCl, 40 mM imidazole, 5% glycerol, 10 mM β-mercaptoethanol, and 1 mM PMSF) and lysed in a C3 homogenizer. After homogenization, cells were centrifuged at 23,000 rpm at 4°C, and the supernatant was loaded onto a 5 mL HisTrap HP Column (Cytiva) equilibrated with Nickel Buffer A (1x PBS, 500 mM NaCl, 40 mM Imidazole, 5% Glycerol, and 10 mM β-mercaptoethanol). The column was washed with Nickel Buffer A to remove non-specificly bound proteins. Exonucleases were eluted over a step-wise gradient to Nickel Buffer B (1x PBS, 500 mM NaCl, 1.5M Imidazole, 5% Glycerol, and 10 mM β-mercaptoethanol). The following fractions were taken for the step wise gradient: 2 × 10 mL fractions of 1% B, 2 × 10 mL 3%, 1 × 10 mL 4%, 1 × 5 mL 5%, 1 × 5 mL 6%, 1 × 5 mL 7%, 1 × 5 mL 8%, 1 × 5 mL 9%, 2 × 10 mL of 10%, 1 × 10 mL 100%. Eluted samples intended for tag-cleavage were digested with Ulp1 protease overnight during dialysis against Dialysis Buffer A (1x PBS, 500 mM NaCl, 10 mM β-mercaptoethanol). Digested protein was loaded onto a 5 mL HisTrap HP Column (Cytiva) to remove cleaved SUMO tag, and the flow-through fraction was dialyzed overnight against Dialysis Buffer B (20 mM Tris-HCl pH 7.5, 300 mM NaCl, and 10 mM β-mercaptoethanol). The dialyzed, tag-free exonucleases were concentrated by ultrafiltration and flash frozen in liquid nitrogen prior to storage at −80°C.

### *In vitro* trimming assay with exonucleases

For RISC assembly, 20 μM of recombinant FLAG-AGO1, 2, 3, and 4 were incubated for 1 hour at 37°C with 2 μM of 5’ end-labeled 23-nt miR-20a, 21-nt let-7a, 22-nt miR-16, or 23-nt miR-19b in 1x Trimming Buffer (25 mM HEPES-KOH pH 7.5, 50 mM KCl, 5 mM DTT, 0.2 mM EDTA, 0.05 mg/mL BSA). The assembled RISC reactions were pulled down through a 2-hour incubation at room temperature with anti-FLAG M2 beads (Sigma Aldrich) that had been pre-washed twice with 1X PBS and 1X Trimming buffer. The beads were washed 8 times with IP wash buffer (50 mM Tris-HCl pH 7.5, 300 mM NaCl, and 0.05% NP-40) and twice with 1x trimming buffer that included either 2 mM MnCl_2_ or MgCl_2_. For the trimming reaction, the beads were incubated with 500 pmol of recombinant exonuclease (ISG20, ISG20 D11N/D94N, TREX1, TREX1 D200N, ERI1, ERI1 D234A, PARN, PARN D28A, or EXO5) for 4 hours at 37°C. The beads were then washed with IP wash buffer 4 times, mixed with 2x urea quenching dye (8 M urea, 1 mM EDTA, 0.05% (w/v) xylene cyanol, 0.05% (w/v) bromophenol blue, 10% (v/v) phenol), and resolved on an 8 M urea 16% (w/v) polyacrylamide gel. Images were analyzed by Typhoon Imaging System (GE Healthcare), quantified by Image Lab (Bio-Rad), and statistically analyzed using GraphPad Prism.

### *In vitro* trimming followed by in vitro cleavage assay with four miRNAs

For RISC assembly, 20 μM recombinant FLAG-AGO3 was incubated with 2 μM of 23-nt miR-20a, 21-nt let-7a, 22-nt miR-16, or 23-nt miR-19b in 1x Trimming Buffer for 1-hour at 37°C. The assembled RISC reactions were pulled down through a 2-hours incubation at room temperature with anti-FLAG M2 beads (Sigma Aldrich) that had bene pre-washed twice with 1X PBS and 1X Trimming buffer. The beads were then washed with IP wash buffer 8 times and twice with a 1x trimming buffer that included 2 mM MnCl_2_. For the trimming reaction, the beads were then incubated with 500 pmol of ISG20 for 4 hours at 37°C. The beads were then washed with IP wash buffer 8-times followed by two washes with 1x Reaction buffer (25 mM HEPES-KOH pH 7.5, 50 mM KCl, 5 mM DTT, 0.2 mM EDTA, 0.05 mg/mL BSA, 0.5 U/μL Ribolock, 5 mM MgCl_2_). For the cleavage reaction, 10 pmol of the spiked target of miR-20a, let-7a, miR-16, or miR-19b was added to the beads with 1X Reaction Buffer and incubated for 1 hour. The beads were mixed with 2X urea quenching dye. The cleavage products were loaded on an 8 M urea 16% (w/v) polyacrylamide gel. Images were analyzed by Typhoon Imaging System (GE Healthcare) and quantified by Image Lab (Bio-Rad).

### Cell culture

HEK293T cells were grown in Dulbecco’s modified Eagle’s medium (DMEM) (Gibco) supplemented with 10% FBS (Gibco) at 37°C in 5% CO2 incubator.

### *In vivo* RNA stability assay

The cells were transfected with 25 nM 5’ end labeled single-stranded 14-nt miR-20a, single-stranded 23-nt miR-20a, 14-nt siRNA-like miR-20a duplex, or 23-nt siRNA-like miR-20a duplex. After a 48-hour transfection, the cells were lysed using 1X RIPA buffer (Cell Signaling Technology) containing 1 mM PMSF and incubated with 400 μg/mL proteinase K at 56°C for 1 hour. Total RNA was extracted using UltraPure™ Phenol:Chloroform:Isoamyl Alcohol (25:24:1, v/v) (Invitrogen™) and precipitated with ethanol. The level of radioactivity of each sample was measured using scintillation counter and the samples adjusted to the same radioactivity were loaded on an 8 M urea 16% (w/v) polyacrylamide gels. Images were analyzed by Typhoon Imaging System (GE Healthcare).

### *In vivo* RNA degradation assay

The cells were co-transfected with 25 nM 5’ end labeled 23-nt siRNA-like miR-20a duplex, and 20 μg of a plasmid which contained either FLAG-MBP or ISG20. After a 48-hour transfection, the cells were lysed using 1X RIPA buffer (Cell Signaling Technology) containing 1 mM PMSF. Total RNA was extracted using UltraPure™ Phenol:Chloroform:Isoamyl Alcohol (25:24:1, v/v) (Invitrogen™) and precipitated with ethanol. The level of radioactivity of each sample was measured using scintillation counter and the samples adjusted to the same radioactivity were loaded on an 8 M urea 16% (w/v) polyacrylamide gels. Images were analyzed by Typhoon Imaging System (GE Healthcare).

### *In vivo* trimming assay with exonucleases

The cells were co-transfected with 25 nM 5’ end labeled 23-nt siRNA-like miR-20a duplex, 10 μg of FLAG-AGO plasmid, and 20 μg of a plasmid which contained either FLAG-MBP or an exonuclease (ISG20, ISG20 D11N/D94N, TREX1, TREX1 D200N, ERI1, ERI1 D234A, PARN, PARN D28A, or EXO5). HEK293T cells were grown in DMEM media (Gibco) supplemented with 10% FBS (Gibco). The cells were co-transfected with 25 nM 5’ end labeled siRNA-like miR-20a duplex, 10 μg of a plasmid of either FLAG-AGO plasmid, and 20 μg of a plasmid of which contained either FLAG-MBP or an either exonuclease (ISG20, ISG20 D11N/D94N, TREX1, TREX1 D200N, ERI1, ERI1 D234A, PARN, PARN D28A, or EXO5). After a 48-hours of the transfection, the cells were lysed using 1X RIPA buffer (Cell Signaling Technology) containing 1 mM PMSF. The cell lysate was incubated with anti-FLAG beads for the immunoprecipitation. The beads were washed 8-times with IP wash buffer (300 mM NaCl, 50 mM Tris-HCl pH 7.5, and 0.05% NP-40) and then mixed with 2X urea quenching dye. The cleavage products were loaded on an 8 M urea 16% (w/v) polyacrylamide gels. Images were analyzed by Typhoon Imaging System (GE Healthcare), quantified by Image Lab (Bio-Rad), and statistically analyzed using GraphPad Prism. The band intensities of tyRNAs (15 nt and shorter) were divided by those of the sum of tyRNAs and the full-length guide RNAs (18 - 23 nt).

### Western blot analysis

The cells were co-transfected with 25 nM 23-nt siRNA-like miR-20a duplex, 10 μg of FLAG-AGO plasmid, and 20 μg of a plasmid which contained either FLAG-MBP or an exonuclease (ISG20, ISG20 D11N/D94N, TREX1, TREX1 D200N, ERI1, ERI1 D234A, PARN, PARN D28A, or EXO5). 48-hours post-transfection, the cells were lysed using 1X RIPA buffer containing 1 mM PMSF. The 280 μg of the whole cell lysate was loaded onto an SDS-PAGE (Bolt™ 4-12% Bis-Tris gel) and transferred to nitrocellulose membrane. The membranes were incubated with primary antibodies: anti-FLAG antibody (1:2,000, Sigma), anti-ISG20 antibody (1 μg/mL, Proteintech), anti-TREX1 antibody (1:1,000, Thermo Fisher Scientific), anti-ERI1 antibody (1:1,000, Proteintech), anti-PARN antibody (1:2,000, FORTIS), anti-EXO5 antibody (0.4 μg/mL, Atlas Antibodies), and/or anti-alpha-tubulin antibody (1:000 μg/mL, Cell Signaling Technology), and secondary antibodies: anti-mouse or rabbit (1:15,000, LI-COR). The protein bands were visualized by Odyssey.

## Acknowledgments

This work was supported by Pelotonia Fellowships (to G.Y.S and M.S.P.), a Center for RNA Fellowship (to G.Y.S.), the NIH (R01GM138997 to K.N.), and the Office of the Director, NIH (S10OD023582).

## Author Contributions

K.N. designed research; G.Y.S., A.C.K., M.S.P., J.S., C.D., H.Z., D.B., N.M., and E.A.E-W. performed experiments and analyzed data. G.Y.S and K.N. wrote the paper with input from the other authors.

## References

1. V. Ambros et al., A uniform system for microRNA annotation. RNA 9, 277–279 (2003).

2. K. Nakanishi, Anatomy of RISC: how do small RNAs and chaperones activate Argonaute proteins? Wiley Interdiscip Rev RNA 7, 637–660 (2016).

3. K. Nakanishi, Anatomy of four human Argonaute proteins. Nucleic Acids Res (2022).

4. K. Nakanishi, Are Argonaute-Associated Tiny RNAs Junk, Inferior miRNAs, or a New Type of Functional RNAs? Front Mol Biosci 8, 795356 (2021).

5. A. M. Stankiewicz, J. Goscik, A. Majewska, A. H. Swiergiel, G. R. Juszczak, The Effect of Acute and Chronic Social Stress on the Hippocampal Transcriptome in Mice. PLoS One 10, e0142195 (2015).

6. S. Deymier, C. Louvat, F. Fiorini, A. Cimarelli, ISG20: an enigmatic antiviral RNase targeting multiple viruses. FEBS Open Bio 12, 1096–1111 (2022).

7. M. F. Thomas et al., Eri1 regulates microRNA homeostasis and mouse lymphocyte development and antiviral function. Blood 120, 130–142 (2012).

8. M. S. Park, G. Sim, A. C. Kehling, K. Nakanishi, Human Argonaute2 and Argonaute3 are catalytically activated by different lengths of guide RNA. Proc Natl Acad Sci U S A 117, 28576–28578 (2020).

9. D. Budinger, S. Barral, A. K. S. Soo, M. A. Kurian, The role of manganese dysregulation in neurological disease: emerging evidence. Lancet Neurol 20, 956–968 (2021).

10. B. A. Friedman et al., Diverse Brain Myeloid Expression Profiles Reveal Distinct Microglial Activation States and Aspects of Alzheimer’s Disease Not Evident in Mouse Models. Cell Rep 22, 832–847 (2018).

11. Y. J. Crow et al., Mutations in the gene encoding the 3’-5’ DNA exonuclease TREX1 cause Aicardi-Goutieres syndrome at the AGS1 locus. Nat Genet 38, 917–920 (2006).

12. M. S. Park et al., Human Argonaute3 has slicer activity. Nucleic Acids Res 45, 11867–11877 (2017).

13. M. S. Park et al., Multidomain Convergence of Argonaute during RISC Assembly Correlates with the Formation of Internal Water Clusters. Mol Cell 75, 725–740 (2019).

